# KIAA1109 is required for survival and for normal development and function of the neuromuscular junction in mice

**DOI:** 10.1101/2022.02.23.481678

**Authors:** Yun Liu, Weichun Lin

**Affiliations:** Department of Neuroscience, UT Southwestern Medical Center, Dallas, TX 75390, USA

**Keywords:** KIAA1109, neuromuscular junction, synaptic transmission, mice

## Abstract

KIAA1109 (4932438A13Rik) is a novel gene linked to Alkuraya-Kucinska Syndrome, an autosomal recessive disorder with severe brain malformations and arthrogryposis in humans. The role of KIAA1109 in mammalian development and function remains poorly understood. Here, we characterize mutant mice deficient in *Kiaa1109* (*Kiaa1109^−/−^*). We report that *Kiaa1109^−/−^* mice died during perinatal stages. These *Kiaa1109^−/−^* embryos exhibited impaired intramuscular nerve growth and reduced sizes of the neuromuscular junction (NMJ) compared with their littermate controls. Electrophysiological analysis further revealed defects in neuromuscular synaptic transmission in *Kiaa1109^−/−^* embryos. Notably, the frequency of spontaneous neurotransmitter release was markedly increased, whereas evoked neurotransmitter release and quantal content were reduced. Furthermore, neuromuscular synapses in *Kiaa1109^−/−^* embryos failed to respond to a repetitive, low frequency stimulation (10Hz). These results demonstrate that KIAA1109 is required for survival in mice and for proper development and function of the NMJ.

**Significance Statement:** This is the first report characterizing the phenotype of mutant mice deficient in KIAA1109 (4932438A13Rik), a novel gene in mammals. We show that KIAA1109 is required for survival in mice and that KIAA1109 plays important roles in normal development and function of the NMJ in mice.

## Introduction

*KIAA1109* was initially cloned from a size-fractionated adult human brain cDNA library (Kikuno et al., 1999) and was later mapped to the chromosome 4q27 region in humans (van Heel et al., 2007). Genetic studies in humans have shown that *KIAA1109* is associated with Alkuraya-Kucinskas Syndrome – an autosomal recessive severe neurodevelopmental disorder characterized by developmental delay, severe brain malformations, arthrogryposis and clubfoot (Filatova et al., 2019; Gueneau et al., 2018; Kumar et al., 2020; Kvarnung et al., 2018; Meszarosova et al., 2020). Based on its sequence, *KIAA1109* encodes for a large protein of 5005 amino acids, and is an evolutionally conserved protein. In Yeast *(Saccharomyces cerevisiae)*, Csf-1, the orthologue of KIAA1109 is predicted as a tube-forming lipid transport protein critical for glycosylphosphatidylinositol anchor biosynthesis in the endoplasmic reticulum (Toulmay et al., 2022). In Zebrafish, *kiaa1109* gene knockdown results in hydrocephaly and abnormally curved notochords in fish embryos reminiscent of clinical features in Alkuraya-Kucinskas Syndrome in human patients (Gueneau et al., 2018). In *Drosophila, Tweek,* an orthologue of KIAA1109 in flies, is involved in regulating membrane endocytosis and synaptic vesicle recycling (Verstreken et al., 2009) and in regulating the size of the NMJs via a phosphatidylinositol 4,5-biophosphate (PI(4,5)P2)- and WSP-dependent pathway (Khuong et al., 2010). In human patient-derived dermal fibroblasts carrying compound heterozygous *KIAA1109* variants, endocytosis and endosome trafficking are also notably compromised (Kane et al., 2019), suggesting the function of KIAA1109 in humans may overlap with the function of its fly ortholog *Tweek*.

Here, we examined the role of KIAA1109 in mouse embryonic development, focusing on the formation and function of the neuromuscular junction (NMJ) in mutant mice deficient in KIAA1109 (*Kiaa1109^−/−^*). We found that ablating KIAA1109 in mice led to perinatal lethality; no *Kiaa1109^−/−^* mice survived beyond postnatal day 0 (P0). Furthermore, *Kiaa1109^−/−^* embryos exhibited marked defects in motor innervation and neurotransmission at the NMJ. The results suggest KIAA1109 plays key roles in neurodevelopment and synaptic neurotransmission at the mouse NMJ.

## Results

### KIAA1109 deficiency leads to perinatal lethality in mice

We obtained heterozygous mice *(Kiaa1109^+/-^*) from the Jackson Laboratory. These *Kiaa1109^+/-^* mice were viable and fertile, with no overt phenotype. To generate *Kiaa1109^−/−^* mice, *Kiaa1109^+/-^* mice were interbred using standard procedures. During our initial analysis, we genotyped a total of 48 mice from 9 litters at weaning stage (after postnatal day 21, P21). These mice were identified as either wild type (+/+) or heterozygote (+/-), no homozygote *Kiaa1109^−/−^* mice were identified after weaning. We therefore turned our attention to newborn pups and we noticed that some newborn pups (5 pups in 6 litters) died shortly after birth. These dead pups were typically smaller than their littermates and were later identified as *Kiaa1109^−/−^* by genotyping. Since respiratory failure is a common cause for neonatal death, we collected lung tissues from pups at P0. We performed a lung floatation assay using a small piece of lung tissue and noticed that the lung tissue from *Kiaa1109^−/−^* pups sank to the bottom in PBS, whereas the lung tissue from *Kiaa1109^+/+^* pups remained afloat (Fig. 1 B), suggesting that the mutant lungs were not inflated by the air. This was further confirmed by histological analysis of the lung sections – the alveoli in the WT were fully expanded whereas the alveoli in the mutant remained unexpanded (Fig.1 C). These observations suggest that *Kiaa1109*^−/−^ mice likely died from respiratory failure.

**Figure 1.**
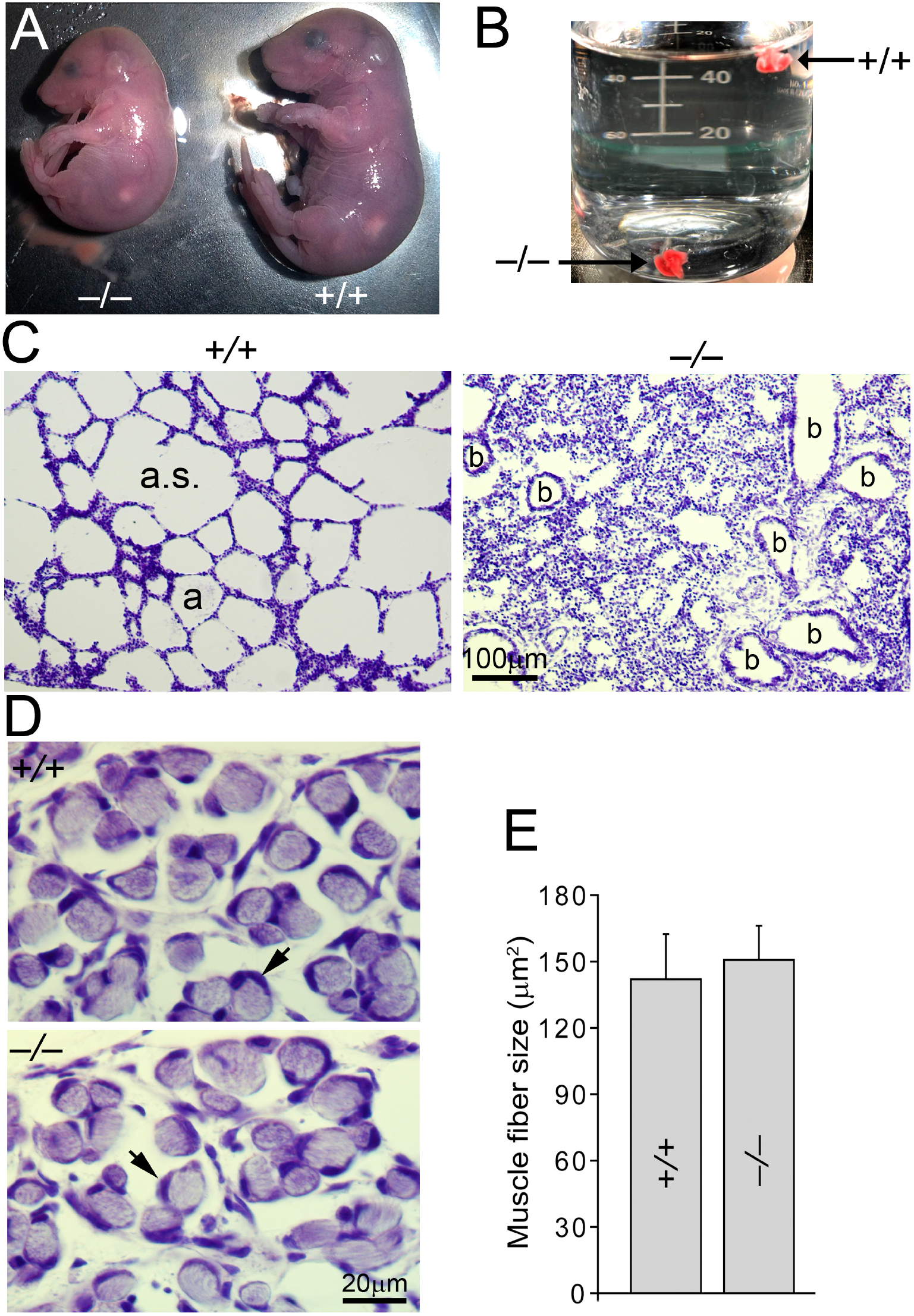
KIAA1109 deficiency leads to perinatal lethality in mice. A: External phenotype of *Kiaa1109^−/−^* and *Kiaa1109^+/+^* embryos at E18.5. Note the mutant embryo (-/-) is smaller and curved (hunchback), compared with its WT littermate (+/+). B: Lung floatation assay: two small pieces of lung tissue (one from WT, and the other from *Kiaa1109^−/−^* at P0) were placed in a beaker filled with PBS. The *Kiaa1109^−/−^* lung tissue sank to the bottom in PBS, whereas the WT lung tissue remained afloat. C: Histological analysis of a lung section (Cresyl Violet staining) shows that the alveoli (a) in WT mouse were fully expanded whereas the alveoli in *Kiaa1109^−/−^* mouse were compressed, indicating breathing failure in the mutant. a: alveoli; b: bronchiole. D and E: Cross sections of cervicis muscles at E18.5 show that muscle fiber sizes were similar between WT (141.94 ± 20.49 μm^2^, N = 3 mice, n = 271 muscle fibers) and *Kiaa1109^−/−^* mice (150.71 ± 15.5 μm^2^, N = 3, n = 265 muscle fibers). Arrowheads in D point to muscle nuclei peripherally localized in both mutant and WT muscle fibers. Scale bars: C: 100 μm; D: 20 μm.

This perinatal lethality precluded further analyses of *Kiaa1109^−/−^* mice at postnatal stages. We therefore focused our subsequent studies on embryonic stages. We collected 107 embryos (at E18.5) from 16 litters (Table 1). Among these embryos, 16 embryos (5 dead, 11 alive) were identified as *Kiaa1109^−/−^* at E18.5. Again, these *Kiaa1109^−/−^* embryos were noticeably smaller than their littermates (Fig. 1 A). In addition, *Kiaa1109^−/−^* embryos typically appeared hunchback with wrist dropping (Fig. 1 A); they exhibited little spontaneous movements but were able to respond to a mild pinch of their skin or tails. Table 1 summarizes total number of embryos at E14.5, E16.5 and E18.5. The numbers in parentheses are the numbers of embryos predicted based on the Mendelian ratio (1:2:1). The observed numbers of *Kiaa1109^−/−^* embryos at E14.5 and at E16.5 were fewer than those predicted, although the differences did not reach statistical significance at these stages (E14.5-E16.5). However, at E18.5, the number of *Kiaa1109^−/−^* mice was significantly reduced (***χ*2** = 7.1245, P = 0.0284). These data indicate that the majority of *Kiaa1109^−/−^* embryos died perinatally and that KIAA1109 was required for survival in mice.

**Table 1.**
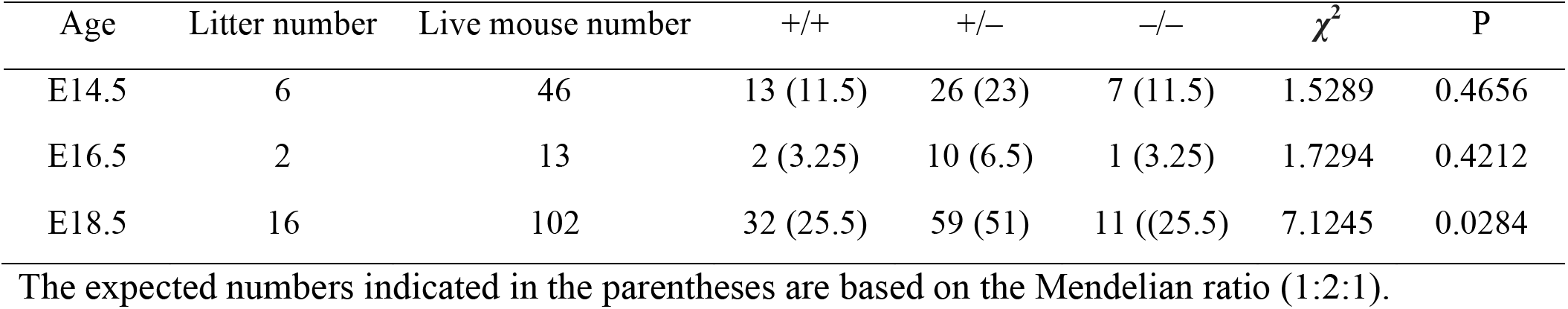
A summary of *Kiaa1109* mouse survival numbers during embryonic stages.

### Abnormal motor innervation in *Kiaa1109^−/−^* mice

The perinatal lethality in *Kiaa1109^−/−^* embryos prompted us to examine the formation and function of the NMJ, as the NMJ is crucial for survival in mice. We analyzed the innervation pattern of embryonic diaphragm muscles by wholemount immunofluorescence staining. Using anti-syntaxin 1 antibodies as a marker for innervation, we detected a significant reduction in the number of nerve branches and innervation area in *Kiaa1109^−/−^* diaphragms, compared with control (Fig. 2 A and C, B and D). Acetylcholine receptors (AChRs) were clustered as an endplate band at the center region of the diaphragm muscles in both *Kiaa1109^−/−^* and control embryos, but the endplate band appeared narrower in *Kiaa1109^−/−^* diaphragms, compared with that of the controls. For example, in the dorsal quadrant of the diaphragm muscle, the average width of the endplate band at E14.5 was 159.27 ± 6.43 μm in *Kiaa1109^−/−^* muscles, comparable to the band width of 181.38 ± 16.99 μm in control muscles (Fig. 3 C). At E18.5, the average width of the endplate band in *Kiaa1109^−/−^* muscles (250.75 ± 10.75 μm, N = 3 mice) was significantly (P = 8.21×10^-5^) less than that of the controls (382.46 ± 8.4 μm, N = 3 mice) (Fig. 3 D).

**Figure 2.**
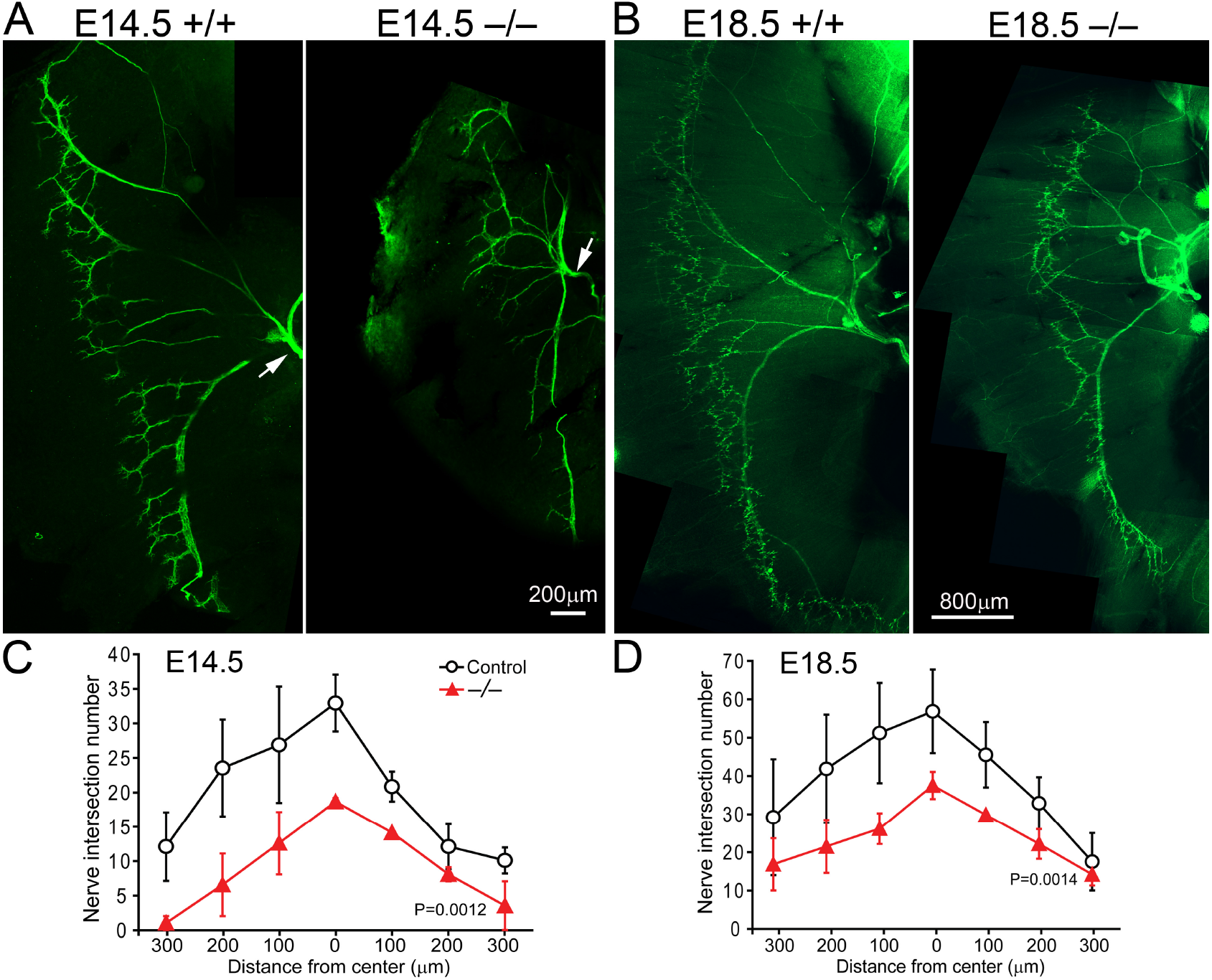
Aberrant innervation pattern in *Kiaa1109^−/−^* muscles. Hemi-diaphragm muscles (right side) at E14.5 (A) and E18.5 (B) were labeled by anti-syntaxin-1 antibodies for the visualization of the phrenic nerves. The innervation territories appear narrower in *Kiaa1109^−/−^* muscles, compared with the WT. C-D: Quantification shows that the numbers of nerve branches in *Kiaa1109^−/−^* embryos are significantly reduced at both E14.5 (P = 0.0012, student paired *t*-test) and E18.5 (P = 0.0014, student paired *t*-test). Scale bars: A, 200 μm; B: 800 μm.

**Figure 3.**
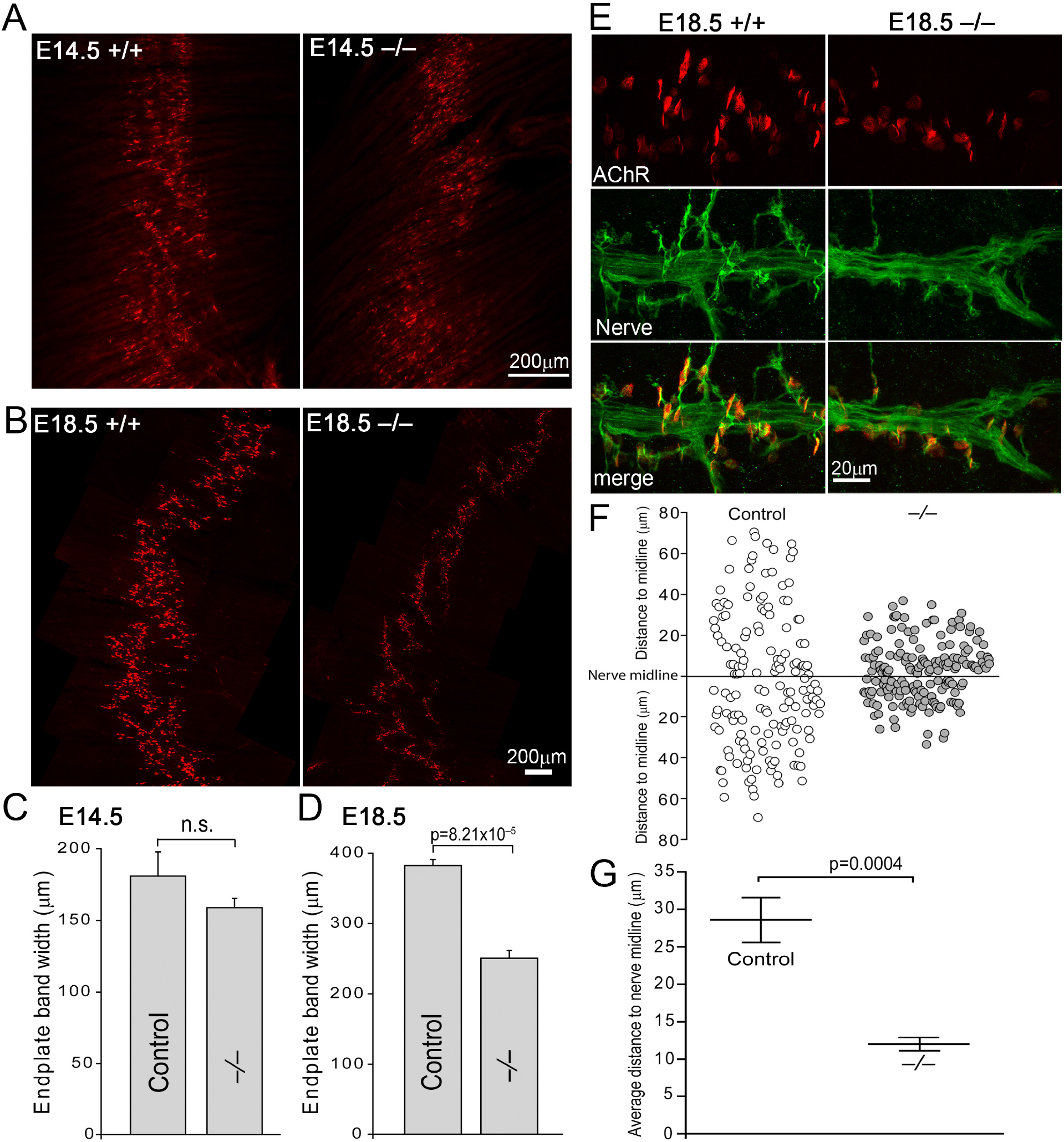
Reduced endplate band in *Kiaa1109^−/−^* muscles. Confocal images of the dorsal right quadrant of the diaphragm muscles labeled by Texas Red conjugated α-bungoratoxin at E14.5 (A) and E18.5 (B). The width of the endplate band is reduced in *Kiaa1109^−/−^* muscles compared to the WT. C-D: Quantification of the width of the endplate band at E14.5 *(Kiaa1109^−/−^*: 159.27 ± 6.43 μm, N = 3 mice; control: 181.38 ± 16.99 μm, N = 4) and E18.5 *(Kiaa1109^−/−^*: 250.75 ± 10.75 μm, N = 3; control: 382.46 ± 8.4 μm, N = 4). E: High-power views of nerves and individual AChR clusters. F: Scatter plots display the distance of the nerve terminals from the midline of the intramuscular nerve trunk. G: Comparison of average distance of the nerve terminals from the nerve trunk in *Kiaa1109^−/−^* (11.97 ± 0.84 μm, N = 3 mice, n = 185 synapses) and the control (28.65 ± 3.26 μm, N = 3 mice, n = 157 synapses). Scale bars: A and B, 200 μm; E: 20 μm.

This narrow distribution of the endplate band in *Kiaa1109^−/−^* muscles appeared consistent with their branching pattern of intramuscular nerves, which extend numerous fine branches (pre-terminal axons) to muscle fibers. Compared to controls, *Kiaa1109^−/−^* muscles exhibited fewer and shorter pre-terminal axons (Fig. 3 E). We measured the distance between nerve terminals and intramuscular nerve trunks from which the nerve terminals sprouted and generated scatter plots (Fig. 3 F). The average distance was significantly (P = 0.0004) reduced in *Kiaa1109^−/−^* (11.97 ± 0.84 μm), compared to that of the control (28.65 ± 3.26 μm) (Fig. 3 G). This result suggests an insufficient nerve sprouting from nerve trunks prior to formation of synapses to muscle fibers in *Kiaa1109^−/−^* mice.

To further analyze the development of the NMJ, we labeled diaphragm muscles with antibodies against synaptotagmin 2 (Syt2), choline transporter (CHT), or acetylcholinesterase (AChE) (Fig. 4). In both control and *Kiaa1109^−/−^* muscles, nerve terminals were intensely labeled with anti-Syt2 or anti-CHT antibodies, and were juxtaposed to postsynaptic AChRs (arrows in Fig. 4 A and B). Similarly, anti-AChE labeling was superimposed AChR clusters in both control and mutant muscles (arrows in Fig. 4 C). However, the sizes of AChR clusters (endplates) in mutants were significantly (P = 0.0451) smaller compared with the control (Fig. 4 D).

**Figure 4.**
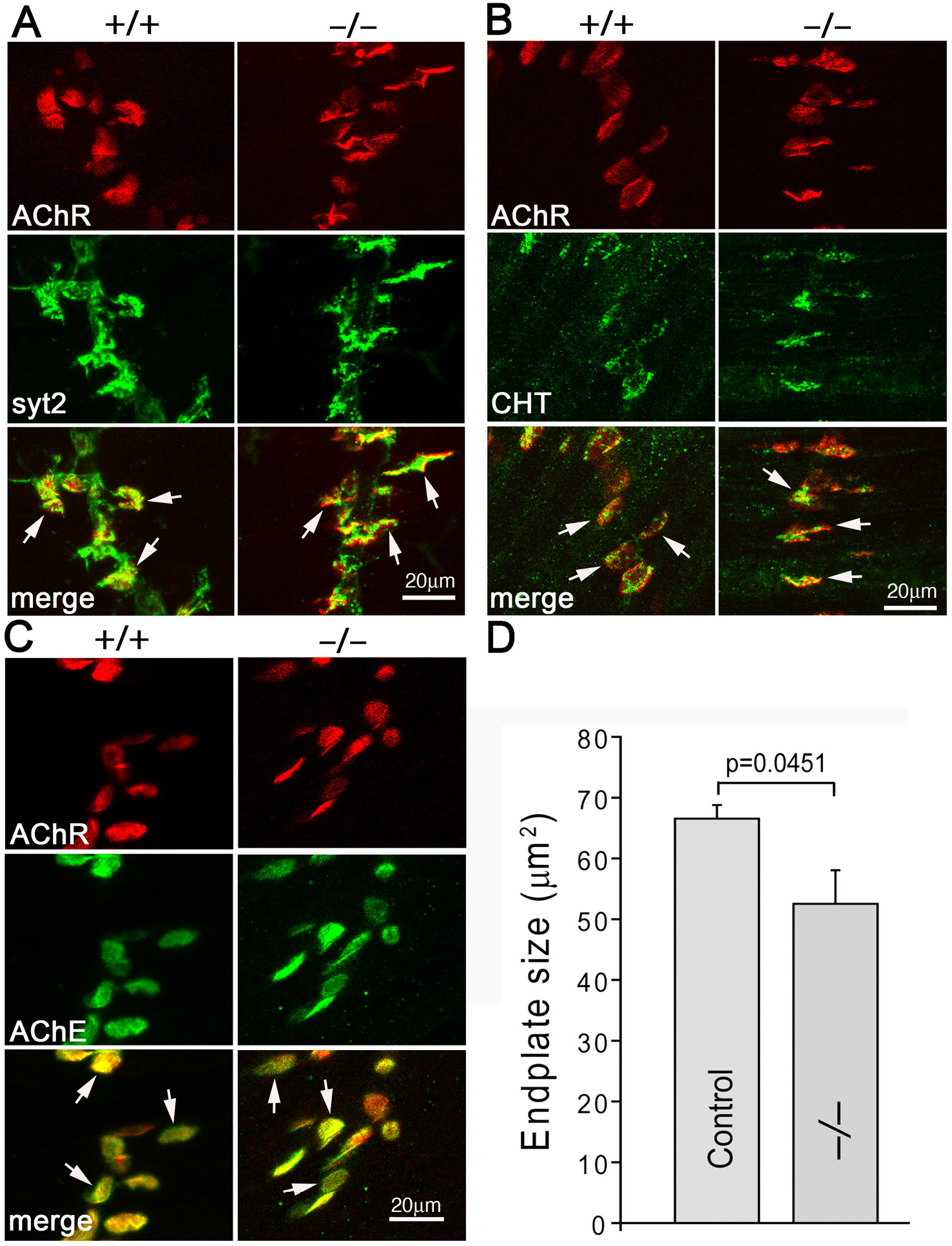
Reduced neuromuscular synapse sizes in diaphragm muscles of E18.5 *Kiaa1109^−/−^* mice. A-C: High power images of the neuromuscular synapses doubly labeled with Texas Red conjugated α-bungoratoxin (red) and anti-synaptotagmin 2 (syt 2) (A), anti-choline transporter (CHT) (B), or antiacetylcholinesterase (AChE) (C). Arrows point to the NMJs. D: Quantification of the size of the endplate. The endplate size was significantly reduced in mutant mice (52.53±5.52 μm^2^, N = 3 mice, n = 481 endplates), compared to that of the control (66.52±2.24 μm^2^, N = 3 mice, n = 363 endplates).

Since defects in muscle development may impair the formation of the neuromuscular junction and retrogradely affect presynaptic nerve growth (Rodriguez Cruz et al., 2020; Sanes and Lichtman, 1999; Tintignac et al., 2015), we examined the skeletal muscle in *Kiaa1109^−/−^* mice. Cross sections of cervicis muscles at E18.5 revealed that muscle fiber sizes in mutants (150.71 ± 15.5 μm^2^, N = 3 mice, n = 265 muscle fibers) were comparable to the WT (141.94 ± 20.49 μm^2^, N = 3 mice, n = 271 muscle fibers) (Fig. 1 E). Muscle nuclei were peripherally localized in both mutant and WT muscles (Fig. 1 D). Overall, no signs of developmental retardation, degeneration or atrophy were detected in the mutant muscles. Taken together, these results show that KIAA1109 deficiency leads to insufficient nerve branching, narrower endplate bands and smaller endplates.

### Increased spontaneous neurotransmitter release in *Kiaa1109^−/−^* mice

To further analyze the NMJ function, we carried out electrophysiology analysis. We isolated phrenic nerve/diaphragm muscle preparation from control and *Kiaa1109^−/−^* mice at E18.5 and recorded synaptic activity using intracellular electrodes. We monitored spontaneous neurotransmitter release by analyzing miniature endplate potentials (mEPPs) at resting membrane potentials. In control muscles, the average frequency of mEPPs was 1.49 ± 0.11 events/min (N = 6 mice, n = 70 cells) (Fig. 5 A and C), consistent with the mEPP frequencies observed in our previous studies (Liu et al., 2009; Liu et al., 2012). However, in *Kiaa1109^−/−^* muscles, a significant (P = 0.0064) increase in mEPP frequency was detected (2.45 ± 0.24 events/min, N = 3 mice, n = 33 cells) (Fig. 5 A and C). The average amplitude of mEPPs in *Kiaa1109^−/−^* mice (2.25 ± 0.37 mV) was similar with that of the control (2.59 ± 0.14 mV) (Fig. 5 B and D), suggesting the quantal size was not affected in *Kiaa1109^−/−^* mice. Furthermore, the parameters of mEPP kinetics, i.e., the rise time and half-width were comparable to those of the control mice (Fig. 5 E and F). These results indicate that KIAA1109 deficiency resulted in an increase in spontaneous transmitter release at the NMJ and suggest that the changes in spontaneous release in mutant NMJs is likely due to a presynaptic alteration.

**Figure 5.**
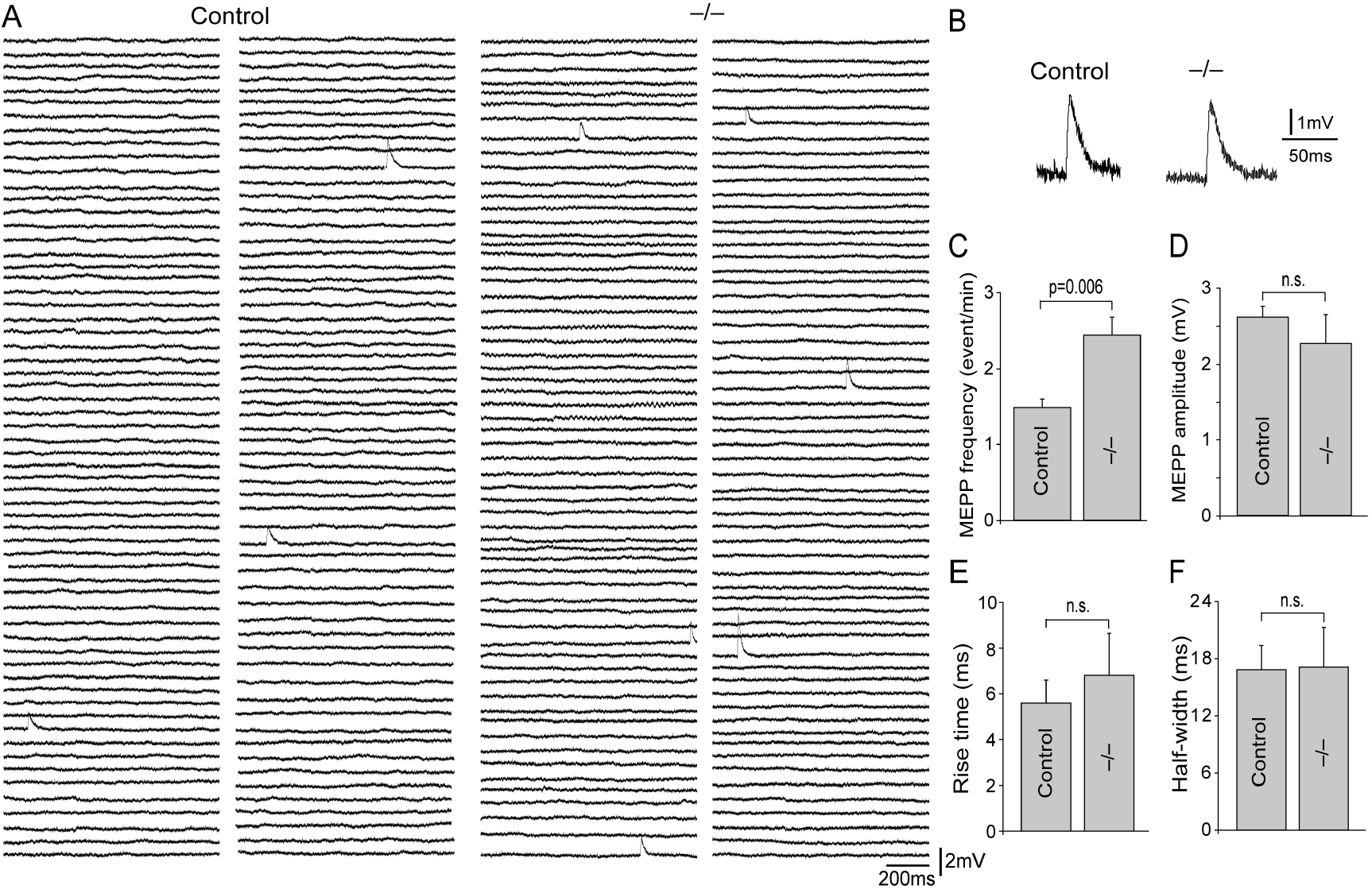
Spontaneous synaptic activity is increased at the NMJs in *Kiaa1109^−/−^* muscles. A: Sample mEPP traces of 2-minute continuous recordings in the diaphragm muscles from the control and *Kiaa1109^−/−^* embryos (E18.5). B: Scale-up examples of mEPPs. C-F: Average mEPP frequency (C), amplitude (D), rise time (E) and half-width (F) are shown in the bar graphs. Notably, the mEPP frequency was increased in *Kiaa1109^−/−^* mice (2.45 ± 0.24 events/min), compared with controls (1.49 ± 0.11 events/min). MEPP amplitudes were comparable between *Kiaa1109^−/−^* (2.25 ± 0.37 mV) and control mice (2.59 ± 0.14 mV). The rise time (E) and half-width (F) are also similar between *Kiaa1109^−/−^* and control (rise time: control, 5.6 ± 1 ms, *Kiaa1109^−/−^*, 6.81 ± 1.84 ms; half-width: control, 16.94 ± 2.55 ms; *Kiaa1109^−/−^*, 17.23 ± 4.19 ms). Numbers of mice (N) and muscle cells (n) analyzed: control, N = 6, n = 70; *Kiaa1109^−/−^*, N = 3, n = 34.

### Defects in evoked synaptic transmission in *Kiaa1109^−/−^* mice

To analyze evoked neurotransmission, we stimulated the phrenic nerve with supra threshold stimuli (2 – 5V, 0.1 ms). In both control and *Kiaa1109^−/−^* mice, muscle action potentials (Fig. 6 C) were readily elicited by nerve stimulation, followed by visually detectable muscle contractions. However, muscle movements in *Kiaa1109^−/−^* mice appeared slower and weaker, compared to those in control mice, nevertheless muscle movement was still sufficient to interfere with stable recordings of evoked responses. We therefore applied 50 μM 3-(N-butylethanimidoyl)-4-hydroxy-2H-chromen-2-one (BHC), a specific myosin inhibitor (Heredia et al., 2016), in Ringer’s solution to suppress muscle movements. In both control and mutant muscles, muscle contraction was markedly suppressed after incubation in BHC for 30 minute – only slight muscle contractions occasionally remained. Upon suppression of muscle contraction, we were able to record stable EPPs, which appeared as either single-peak EPPs (Fig. 6 A) or complex EPPs (Fig. 6 B) (Redfern, 1970). To measure the parameters of EPP amplitude and kinetics, we focused our analyses on the traces with single-peak EPPs (Fig. 6 A). The average amplitude of EPPs in controls was 14 ± 0.66 mV (N = 6 mice, n = 50 cells); whereas the average amplitude of EPPs in *Kiaa1109^−/−^* mice was significantly (P = 0.0018) reduced by 45% to 7.74 ± 1.75 mV (N = 3 mice, n = 38 cells) (Fig. 6 D). The quantal content of *Kiaa1109^−/−^* mice was 4.08 ± 0.91, significantly (P = 0.0176) less than that of the control mice (6.74 ± 0.46) (Fig. 6 E). The rise time and half-width of EPP were not affected in *Kiaa1109^−/−^* mice (Fig. 6 F and G). These results indicate that the evoked neurotransmitter release was compromised in *Kiaa1109^−/−^* mice.

**Figure 6.**
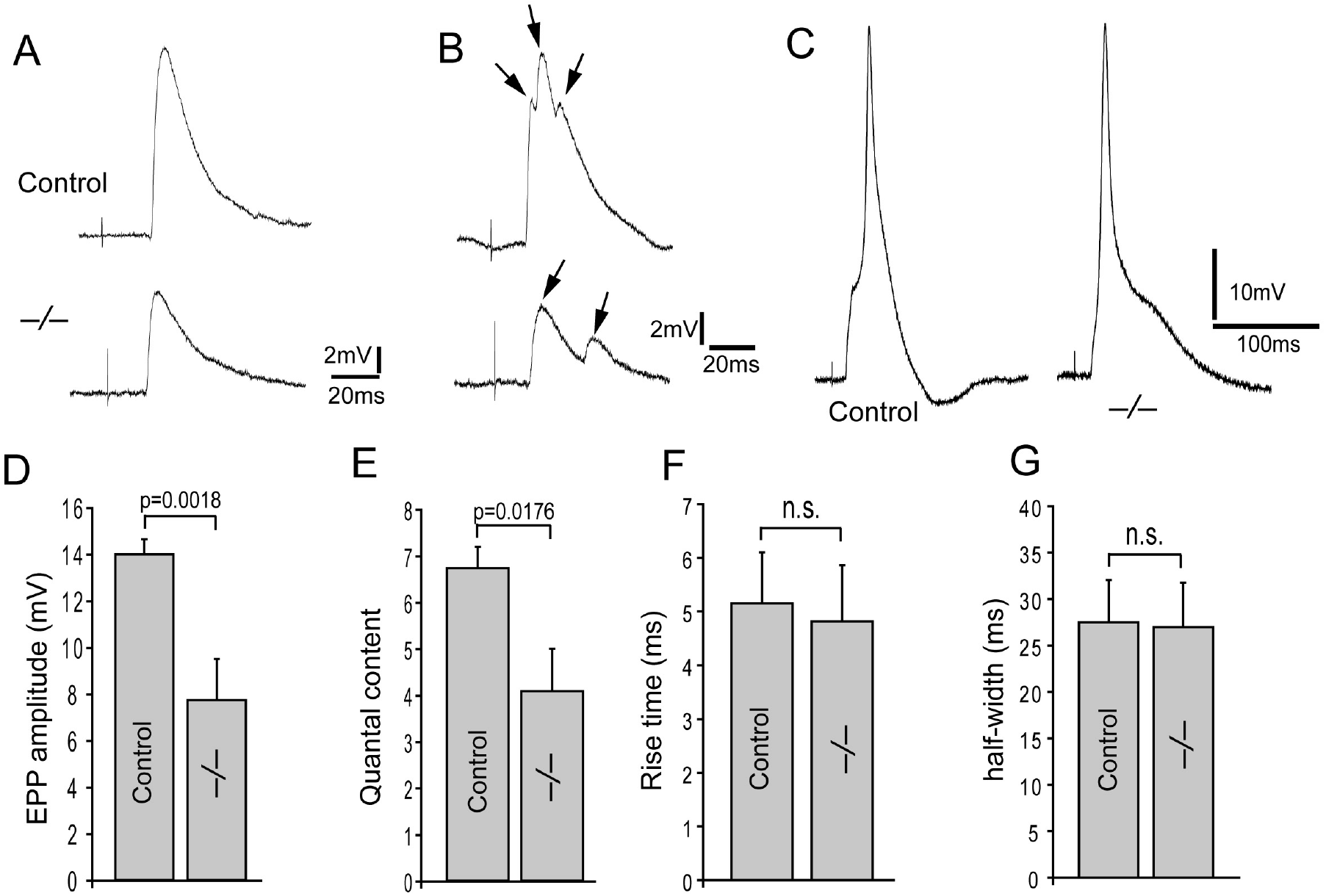
Evoked neurotransmitter release is reduced at the NMJs in *Kiaa1109^−/−^* muscles. A-B: Representative traces of endplate potentials (EPPs) recorded from E18.5 diaphragm muscles. Arrows in B point to EPPs with multiple peaks (complex EPPs). C: Examples of muscle action potentials in response to nerve stimulation. D: EPP amplitude is significantly reduced in *Kiaa1109^−/−^* (7.74 ± 1.75 mV), compared with controls (14 ± 0.66 mV). E: Quantal content is also significantly reduced in *Kiaa1109^−/−^* mice (4.08 ± 0.91) compared with control (6.74 ± 0.46). F-G: Quantification of rise time (F) and half-width (G) between *Kiaa1109^−/−^* and controls (rise time: control, 5.16 ± 0.94 ms; *Kiaa1109^−/−^*, 4.82 ± 1.05 ms; halfwidth: control, 27.52 ± 4.5 ms; *Kiaa1109^−/−^*, 26.97 ± 4.76 ms). Numbers of mice (N) and muscle cells (n) analyzed: control, N = 6, n = 50; *Kiaa1109^−/−^*, N = 3, n = 38.

To further examine neuromuscular function, we stimulated the phrenic nerves with low frequency stimulation. We applied 1 second train stimulation at10 Hz (5 repeats at interval of 6 seconds). In control mice, 97.9% of the cells responded faithfully to each stimulus with an EPP; in contrast, in *Kiaa1109^−/−^* mice, only 54.7% of the cells responded to each stimulus, whereas 45.3% of the cells exhibited at least 1 or more (up to 18) neurotransmission failures (Figure 7). This result indicates that neuromuscular synapses in *Kiaa1109^−/−^* embryos failed to respond to a repetitive, low frequency stimulation (10 Hz). Together, our electrophysiological analyses demonstrate that the NMJ function is markedly compromised in *Kiaa1109^−/−^* mice.

**Figure 7.**
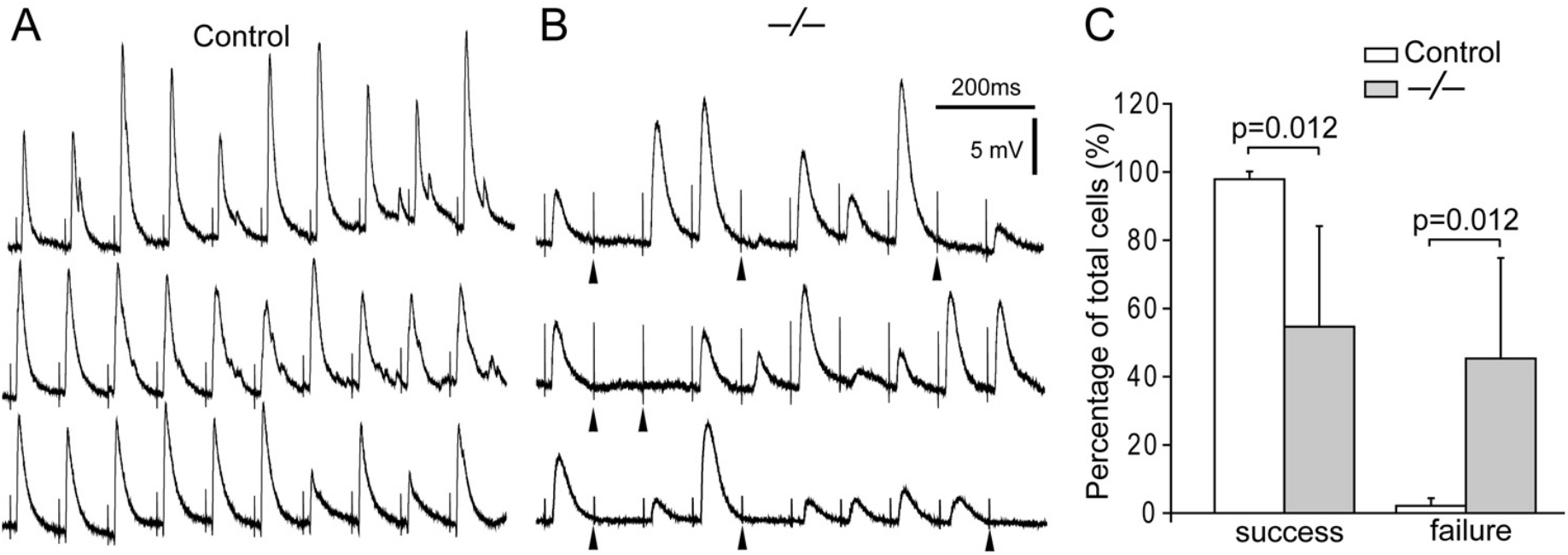
Intermittent synaptic transmission failure in *Kiaa1109^−/−^* muscles. A-B: Sample traces of EPPs in diaphragm muscle (E18.5) in response to 1-second train of supra-threshold stimuli (10 Hz) applied to the phrenic nerve. Such a train of stimuli evoked a train of 10 EPPs in control embryos (A). In contrast, the NMJs in *Kiaa1109^−/−^* diaphragm muscles exhibited intermittent failures of EPP in response to the 10 Hz stimulation as indicated by arrows beneath stimulus artifacts. C: Quantification of the percentages of cells with success or failures in response to the repetitive stimuli. In controls, 97.9% of the cells responded to every stimulus (N = 8 mice, n = 85 cells). In contrast, only 54.7% of the mutant end plates responded to every stimuli, and 45.3% exhibited intermittent synaptic transmission failures (N = 3 mice, n = 38 cells).

## Discussion

In this study, we report the developmental defects in mutant mice deficient in Kiaa1109. We show that the loss of KIAA1109 leads to perinatal lethality in mice, demonstrating that *KIAA1109* is essential for embryonic development and survival. This is consistent with previously reports on *Tweek* mutants in *flies* (Verstreken et al., 2009) and on KIAA1109 variants in human that are associated with Alkuraya-Kucinskas Syndrome (Filatova et al., 2019; Gueneau et al., 2018). The observed phenotype in *Kiaa1109^−/−^* mice – reduced nerve branching and narrower innervation territory – suggest that KIAA1109 is involved in nerve outgrowth during development, as membrane trafficking and addition to the tip of growth cones are key events that lead to axon outgrowth and elongation and synapse formation (Bloom and Morgan, 2011; Futerman and Banker, 1996; Pfenninger, 2009; Rodemer et al., 2020). Similarly, fibroblasts obtained from patients with compound heterozygous *KIAA1109* variants show abnormal endocytosis and endosomal recycling in patient fibroblasts, (Kane et al., 2019). Together, these studies suggest that KIAA1109 plays an evolutionally conserved role in membrane trafficking.

Our electrophysiological studies reveal marked defects in neuromuscular synaptic transmission in *Kiaa1109^−/−^* mice. These defects appear primarily as pre-synaptic in nature as demonstrated by the significant increase in spontaneous transmitter release (mEPP frequency) and decreased evoked transmitter release (EPP amplitude and quantal content). These phenotypes share a common feature with mutant mice deficient in synaptic vesicle proteins such as synaptobrevin (Liu et al., 2011; Liu et al., 2019), SNAP25 (Washbourne et al., 2002), synaptotagmin 2 (Pang et al., 2006), and syntaxin 1B (Wu et al., 2015). Since both spontaneous and evoked neurotransmission require the fusion of synaptic vesicles with the pre-synaptic membrane which leads to exocytosis and neurotransmitter release, the defects in neuromuscular synaptic transmission in *KIAA1109* mutant mice suggest that KIAA1109 may also be involved, albeit indirectly, in vesicle exocytosis.

More importantly, the intermittent transmission failures in mutant NMJs in response to repetitive stimulation suggest that, in the absence of KIAA1109, neuromuscular synaptic transmission cannot be maintained at normal levels. This finding is consistent with the phenotype observed in *Tweek* mutants in Drosophila (Verstreken et al., 2009). The failure to maintain normal levels of neurotransmission could result from multiple factors including a smaller synaptic vesicle pool, compromised vesicle mobilization and defective membrane trafficking and recycling. Indeed, synaptic vesicle numbers at the NMJ and neuromuscular endocytosis are reduced in *Tweek* mutant flies (Verstreken et al., 2009). In addition, defective cargo sorting and abnormal endosomal trafficking have been reported in fibroblasts derived from patients with *KIAA1109* mutations (Kane et al., 2019). Since endosome trafficking is crucial for presynaptic membrane retrieval and vesicle recycling (Buckley et al., 2000; Chanaday et al., 2019; Morgan et al., 2013; Sudhof, 2000), it is plausible that KIAA1109 functions in mammals in a manner similar to Tweek in *Drosophila.* Nevertheless, the mechanisms whereby KIAA1109 regulates synaptic transmission require further investigation.

## Methods and materials

### Mice

*Kiaa1109* mutant mice [B6N(Cg)-4932438A13Rik^tm1b(EUCOMM)Hmgu^/J, Stock No.026878] were generated by the Knockout Mouse Project (KOMP) at The Jackson Laboratory (Bar Harbor, Maine, USA) using embryonic stem cells provided by the International Knockout Mouse **C**onsortium. We obtained heterozygotes *(Kiaa1109^+/−^)* from the Jackson Laboratory; these *Kiaa1109^+/−^* mice were viable and fertile. We generated *Kiaa1109^−/−^* mice from interbreeding of *Kiaa1109^+/−^* mice by timed mating. The day when a vaginal plug first appeared was designated as embryonic (E) day 0.5. A total of 161 mouse embryos (E14.5-E18.5) were collected by Cesarean section of anesthetized pregnant female mice. Genotyping was carried out by PCR using the following primers: wild-type allele forward GGG ATA TGG CAG AGA AGC TG, reverse AAA ACA ATT GGC TTA GAG ACT TCA; mutant allele forward CGG TCG CTA CCA TTA CCA GT, reverse GAC CAC ACA AAT CCC TTG GT.

All experimental protocols followed National Institutes of Health (NIH) guidelines and were approved by the University of Texas Southwestern Institutional Animal Care and Use Committee.

### Histology staining

The cervical portions of whole embryos (E18.5) and lung tissues collected from P0 mice were fixed with 2% paraformaldehyde (PFA) at 4°C overnight. Samples were washed with PBS, equilibrated with 30% sucrose and mounted in OCT compound. Samples were then transversely cryosectioned at a thickness of 12 μm and sections were stained with 0.5% Cresyl Violet. Images were acquired on an Olympus BX51 upright microscope with Nomarski optics.

### Whole mount immunofluorescence staining

Immunofluorescence staining was carried out on whole mount muscles as previously described (Liu et al., 2008). Diaphragm muscles were dissected from mouse embryos (E14.5 – E18.5). Muscle samples were fixed in 2% paraformaldehyde (PFA) at 4°C overnight. Then samples were extensively washed with PBS and incubated with Texas-Red conjugated α-bungarotoxin (α-bgt) (2 nM, Molecular Probes) at room temperature for 30 minutes. Samples were then incubated with primary antibodies at 4°C overnight. Primary antibodies used were: syntaxin-1 (I379), synaptotagmin-2 (I735) (1:1000, gifts from Dr. Thomas Südhof, Stanford University School of Medicine, Palo Alto, CA) (Pang et al., 2006), choline transporter (1:1000, gift from Dr. Randy Blakely, Vanderbilt University School of Medicine, Nashville, TN) (Ferguson et al., 2004) and acetylcholinesterase (1:1000, gifts from Dr. Palmer Taylor, Skaggs School of Pharmacy & Pharmaceutical Sciences, UC San Diego, CA, USA). After three washes with PBS samples were then incubated with fluorescein isothiocyanate-conjugated (FITC) secondary antibodies (1:600) at 4°C overnight. Samples were washed in PBS and mounted in Vectashield mounting medium (H-1000, Vector Laboratories, Inc., Burlingame, CA, USA). Fluorescence images were acquired with a Zeiss LSM 880 confocal microscope.

### Morphometric analysis

High power images (40×, N.A. 0.75) collected on Cresyl Violet-stained cross sections of whole embryos at cervical level were used for measurement of muscle fiber size. Cervicis muscles on the sections were distinguished according to the atlas of mouse embryos. Cross areas of individual muscle fibers were manually traced and areas were measured by using Image J software. Nerve branching was quantified using the methods describe previously (Kaplan et al., 2018; Liu et al., 2019). Confocal images were captured on the right hemi-diaphragms of E14.5 and E18.5 embryos. A vertical straight line perpendicular to the long axis of the muscle was drawn at the center of the endplate band. Then, three straight lines in parallel with the first line were respectively drawn to the left and right sides of the first line at 100 μm interval. The numbers of intersections between nerve branches and the lines were manually counted. The width of the endplate band was measured from the right dorsal quadrant of E14.5 and E18.5 diaphragms using NIH Image J. First, the endplate band labeled with Texas-Red conjugated α-bgt was manually contoured. Then, 50 straight lines in parallel with the long axis of muscle fibred were drawn evenly within the contoured area. The length of each straight line was measured and the average length of the 50 lines was calculated as the width of the endplate band. The sizes of individual endplates were measured from high power confocal images (63 ×, N.A.1.4 oil) by using NIH Image J. The distance of individual synapses to the intramuscular nerve trunk was measure based on high power confocal images (63 ×, N.A. 1.4 oil) obtained from the right ventral region of the diaphragm muscles (E18.5). A straight line in parallel with the long axis of the nerve trunk was drawn at the central of the nerve and designated as the midline. Then a vertical straight line from the center of the endplate to the midline was drawn and the length of the line was measured as the distance of the synapse to the nerve trunk.

### Electrophysiology

Intracellular recordings were carried out on diaphragm muscles from *Kiaa1109^−/−^* and littermate control (both *Kiaa1109^+/+^* and *Kiaa1109^+/–^)* mice at E18.5 as previously described (Liu et al., 2008). Acutely isolated phrenic nerve-diaphragm muscles were mounted on a Sylgard coated dish and bathed in oxygenated (95% O_2_, 5% CO_2_) Ringer’s solution (136.8 mM NaCl, 5 mM KCl, 12 mM NaHCO_3_, 1 mM NaH_2_PO_4_, 1 mM MgCl_2_, 2 mM CaCl_2_, and 11 mM d-glucose, pH 7.3). The muscle samples were then incubated in Ringer’s solution containing 50 μM 3-(N-butylethanimidoyl)-4-hydroxy-2H-chromen-2-one (BHC), a specific myosin inhibitor (Heredia et al., 2016), for 30 minutes prior to the recording in the same solution. To evoke synaptic transmission, the phrenic nerve was stimulated with supra threshold stimuli (2 – 5V, 0.1 ms) via a glass suction electrode connected to an extracellular stimulator (SD9, Grass-Telefactor, West Warwick, RI). Endplate regions were identified using a water-immersion objective (Olympus BX51WI). Muscle membranes were penetrated with sharp microelectrodes (resistance 20–40 MΩ) filled with 2 M potassium citrate and 10 mM potassium chloride. Muscle membrane potentials were acquired with an intracellular amplifier (AxoClamp-2B) and digitized with Digidata 1332A (Molecular Devices, Sunnyvale, CA, USA). MEPPs were analyzed with Mini Analysis Program (Synaptosoft, Inc., Decatur, GA) and EPPs were analyzed with pClamp 10.7 (Molecular Devices, CA). The rise time of mEPPs or EPPs was calculated as the time taken by the membrane potential to rise from 10% to 90% of the peak values of mEPPs or EPPs. Quantal content, defined as the number of acetylcholine quanta released in response to a single nerve impulse, was estimated by using a direct method: dividing the mean amplitude of EPPs by the mean amplitude of mEPPs of the same cell (Boyd and Martin, 1956; Hubbard et al., 1969; Wood and Slater, 2001).

### Data analysis

Data were presented as mean ± standard error of the mean (SEM). Statistical analyses were carried out using SigmaPlot or Microsoft Excel software. Chi-square test was used to analyze the survival of mice. For quantitative morphometric analyses, student paired *t*-test was used to compared nerve branching between mutant and control groups. Student *t*-test was used to determine the difference of endplate band width, endplate size and muscle fiber size between mutant and control groups. For electrophysiological analyses, the mEPP frequency, amplitude, mEPP rise time (10% ~ 90%), half width and EPP amplitude, quantal content, EPP rise time (10% ~ 90%) and half width were compared between mutant and control groups by using student *t*-test. The difference between mutant and control groups is considered statistically significant if the P-value is less than 0.05.

## Acknowledgments

We would like to thank Ms Qiaohong Ye for her excellent technical assistant, and Drs. Beverly Rothermel, Jane Johnson and Joseph McArdle for their comments on earlier drafts of the manuscript. This work was supported by grants from NIH/NINDS (R01 NS055028), the Edward Mallinckrodt, Jr. Foundation, and the Cain Foundation in Medical Research.

## Notes

### Competing Interest Statement

The authors have declared no competing interest.

